# CrispRdesignR: A Versatile Guide RNA Design Package in R for CRISPR/Cas9 Applications

**DOI:** 10.1101/805630

**Authors:** Dylan Beeber, Frédéric JJ Chain

## Abstract

The success of CRISPR/Cas9 gene editing applications relies on the efficiency of the single guide RNA (sgRNA) used in conjunction with the Cas9 protein. Current sgRNA design software vary in the details they provide on sgRNA sequence efficiency and are almost exclusively restricted to model organisms. The *crispRdesignR* package aims to address these limitations by providing comprehensive sequence features of the generated sgRNAs in a single program, which allows users to predict sgRNA efficiency and design sgRNA sequences for systems that currently do not have optimized efficiency scoring methods. *crispRdesignR* reports extensive information on all designed sgRNA sequences with robust off-target calling and annotation and can be run in a user-friendly graphical interface. The *crispRdesignR* package is implemented in R and has fully editable code for specialized purposes including sgRNA design in user-provided genomes. The package is platform independent and extendable, with its source code and documentation freely available at https://github.com/dylanbeeber/crispRdesignR.

## Introduction

The CRISPR/Cas9 system has attracted attention in recent years for its ability to edit and regulate DNA in a wide variety of organisms and cell types. Using a strand of single guide RNA (sgRNA), the Cas9 protein is able to search a cellular genome and induce double stranded breaks at a target sequence complementary to the sgRNA that can then be modified^1^. However, several sequence features of the sgRNA and surrounding DNA sequence can influence the enzymatic activity of Cas9^2^. Crucially, the genomic DNA must contain a protospacer adjacent motif (PAM) in the region immediately following the 3’ end of the target DNA for Cas9 to recognize the sequence^1^. Other sgRNA sequence features like nucleotide composition, presence of homopolymers, and self-complementarity can affect the activity of the sgRNA^2^.

The efficiency of the sgRNA is a major factor in the success of Cas9 gene editing applications^2^. To predict the efficiency of sgRNA sequences, scoring methods have been developed by applying machine learning techniques to CRISPR/Cas9 experimental data^3,4,5^. These efficiency scoring methods are accurate within the parameters of the experiments they were based on. However, the predictions are not necessarily generalizable to Cas9 applications in all cell types, organisms, and PAMs not included in the efficiency scoring experimental data. At their most predictive, scoring methods have been shown to only explain about 40% of the variation in efficiency for most guides^6^. Known sequence features that decrease sgRNA efficiency are not always considered by scoring models^3,4^, which could result in suggesting inactive sgRNAs. The predictive power of these machine learning models may be improved by considering their predictions along with the known effects of sequence features in the genome.

Potential sgRNA sequences that contain a sequence feature not conducive to Cas9 enzymatic activity can be scored highly by efficiency scoring methods that have not been trained on that feature. In order to generate the most active sgRNA, sequence features must be considered alongside efficiency scoring, however current programs designed to identify suitable sgRNAs often do not report all sequence features relevant to sgRNA efficiency. This forces users to run multiple programs to obtain all pertinent information. Features like sgRNA self-complementarity, presence of homopolymers, and potential off-target effects can drastically affect experimental outcomes and are often not considered by scoring models^3,4^. sgRNA sequences that are able to form hairpins with themselves or with other regions of the RNA backbone have been shown to either reduce or increase activity in separate situations^7,8^. Homopolymers that contain 4 or more consecutive identical base pairs (e.g. GGGG) can decrease cutting activity, and a homopolymer with 4 consecutive T’s will be terminated prematurely in systems that utilize RNA polymerase III to create the sgRNA^7^. It is possible for Cas9 to target and cleave DNA sequences with multiple mismatches to the guide RNA resulting in off-target effects^3^. While often problematic for those working with Cas9, these off-target sequences as well as hairpins and homopolymers can be predicted from the sequence features of the guide RNA. Such features are expected to affect activity more consistently across different cell types, organisms and PAMs than specific nucleotide position features^2^.

We have developed the R package *crispRdesignR* to improve upon current sgRNA design software for CRISPR/Cas9 applications by providing all guides that match a customizable PAM sequence within a target region of any genome using the advanced Doench Rule Set 2 predictive model^3^, and by reporting sequence features often missing from other available programs but important in the CRISPR/Cas9 system including the GC content, self-complementarity, presence of homopolymers, and potential off-target effects for each candidate sgRNA. This is especially useful for working with non-standard Cas9 applications where the efficiency score may not be reliable. An optional table can be generated that displays supplementary information on where the potential off-target effects occur in a user-selected genome. The *crispRdesignR* package can also be utilized with a graphical user interface for easier accessibility to non-bioinformaticians. In addition, the flexibility of this R package allows users to design sgRNAs in non-model organisms by inputting custom genomes and annotation files for analysis, highlighting the versatility of *crispRdesignR*.

## Materials and Methods

### Model Features

The predictive sgRNA efficiency scoring model used in *crispRdesignR* examines the same features as the Doench model^3^ except for the cut site within the resulting protein, because not every Cas9 target site is located in a protein encoding region. Our program employs a gradient boosted regression model trained on the FC and RES data set used in Doench Rule Set 2. The FC and RES data sets^3^ contain about 5000 sgRNA sites plus context sequence (30-mer) for a variety of different genes. Ranks for each sgRNA site are calculated from read counts and normalized between 0 and 1, which is used by the gradient boosting algorithm gbm^15^ to predict sgRNA activity. The Doench 2016 scoring method is trained on guide RNA utilizing the 5’NGG3’ PAM sequence. When designing guides for custom PAM sequences, crispRdesignR does not change the scoring method as many of the sequence features considered by Doench 2016^3^ are unrelated to the PAM sequence. It is however important to note that the accuracy score provided is expected to be less accurate when designing sgRNA sequences with custom PAMs.

The presence of specific nucleotides at certain positions in an sgRNA target site can influence the activity of that site. crispRdesignR will consider the single and dinucleotides at each position and convert them into features that our machine learning model uses to predict activity. In accordance with the Doench Rule Set 2^3^, our model accounts for the presence of position-dependent single nucleotides, position-dependent dinucleotides, single nucleotide count, dinucleotide count, GC count, nucleotides that bookend the PAM sequence, and thermodynamic features of the target sequence plus context region (30-mer). As in Doench Rule Set 2, nucleotide features are one-hot encoded, meaning that the presence of a nucleotide in a position is either “off” (0) or “on” (1). This leads to four features for each single nucleotide position (A, C, T, or G) and sixteen features for each dinucleotide position (AA, AC, AG, AT, etc.). One-hot encoding of these features is crucial for accurate machine learning predictions and is made possible by the vtreat package^9^. A position-independent total count of single and dinucleotides is also used. This is simply the number of each specific nucleotide and dinucleotide combination in the 30-mer. Four features counting each single nucleotide and sixteen features counting each dinucleotide are recorded.

The GC count of the target site (20-mer) is taken and converted into a single feature (a number between 0 and 20). However, two additional GC features are taken, one binary variable for if the GC count is above 10 and another for if the GC count is below 10. The two nucleotides that bookend the “GG” of the PAM site are one-hot encoded as a dinucleotide feature. These are the nucleotides at position 25 and 28 of the 30-mer. As with the position-dependent dinucleotide features, these two nucleotides are converted into 16 binary features, one for each possible dinucleotide combination.

Four thermodynamic features are recorded, one for the predicted melting temperature (Tm) of the sgRNA plus context sequence (30-mer), one for the Tm of the five nucleotides upstream from the PAM (positions 20-24), one for the Tm of the eight nucleotides upstream from the previous 5-mer (positions 12-19) and one for the Tm of the five nucleotides upstream from the 8-mer (positions 7-11). The Doench Rule Set 2 uses the Tm_staluc function from biopython to calculate the Tm of these regions, so the function employed by *crispRdesignR* mirrors the Tm_staluc function using thermodynamic data from Allawi and SantaLucia^10^.

### Model Predictions

The model features were used to train a gradient boosted regression model with the R package gbm^11^ on the FC and RES data used by the Doench Rule Set 2. Position-dependent features that contained no variation due to the restrictive PAM site were removed. Other features that showed no impact on the predictive power of the model were also removed. To predict the efficiency of package-generated sgRNA target sequences, the same features collected to design the model are collected for each possible target site. The generated data are then run through the gbm package and return a number from 0 to 1 for each target site, with 0 indicating less activity and 1 indicating greater activity.

### Off-Target Annotation

Users may search any genome that is provided through the BSgenome package^12^. BSgenome also allows users to import custom genomes and DNA sequences from FASTA files (using the *forgeBSgenomeDataPkg* command on a seed file that describes the paths to the raw sequence data in FASTA format; more information can be found in the BSgenome documentation). Genome annotation files (.gtf) can be acquired through the Ensembl and BioMart databases or users can upload their own. Larger genomes should be loaded as a compressed .gtf file (.gtf.gz) due to size limitations.

When off-target searching is on, each sgRNA sequence is checked for the presence of possible off-target sequences with up to four mismatches in the 20-mer. Off-target sequences must match the rules of the PAM site or be included in the list of possible 5’NGG3’ PAM mismatches made available by Doench *et al*.^3^. Off-target sequences that contain 4 mismatches and do not directly match the PAM sequence are not reported by crispRdesignR as they are highly unlikely to be active^3^. The *matchPattern*() function available in the package BioStrings^13^ is used to collect data on each possible off-target sequence. *matchPattern*() searches the target genome for matching patterns with between 1 and 4 mismatches. Indels are not considered when searching for matches. When searching genomes with many base pairs (e.g. over 1 billion) it is recommended to keep the DNA query sequence under 500 base pairs to keep the search time to several minutes. While the *matchPattern*() function is slower than other match finding methods because it does not require the genome to be pre-indexed, which itself takes additional time, this method allows users to easily search uploaded custom genomes without prior processing.

The locations of the possible off-target sequences are cross referenced with a user supplied genome annotation file (.gtf) and reports an off-target information table listing each possible off-target along with the sgRNA target site that it matches. *crispRdesignR* reports sgRNA target sequences and other perfect genomic matches in the off-target annotation table so that the user may verify their target location within the genome. The off-target information table lists the sequence type of the off-target, as well as the gene ID, gene name, and exon number. A cutting frequency determinant (CFD) score for each off-target is also listed in the off-target annotation table, which is calculated using data from Doench *et al*.^3^ to estimate the likelihood of Cas9 targeting this sequence. Each mismatch position is assigned a value based on the change from one specific nucleotide to another and the values are multiplied, producing a number between 1 and 0, with 1 being more likely to be targeted and zero being less likely. crispRdesignR does not consider the position of the query target DNA sequence when finding possible off-targets so that the user may verify the location of their sgRNA target sequences within the genome in the off-target annotation table.

### Functions

All data is generated with a single function in R^13^: sgRNA_design(*userseq, genomename, gtfname, userPAM, calloffs* = TRUE, *annotateoffs* = TRUE).

- *userseq*: The target sequence with which to generate sgRNA guides. Can either be a character sequence containing DNA bases (A,C,T,G) or the name of a FASTA or text file in the working directory.
- *genomename*: The name of a genome (in BSgenome format) to check for off-targets and provide locations for sgRNA guides. These genomes can be downloaded through BSgenome or compiled by the user.
- *gtfname*: The name of a genome annotation file (.gtf) in the working directory to annotate sgRNAs and off-target sequences.
- *userPAM*: An optional argument used to set a custom PAM for the sgRNA. If not set, the function will default to the “NGG” PAM. Warning: the accuracy of Doench efficiency scores has only been tested for the “NGG” PAM.
- *calloffs*: If TRUE, the function will search for off-targets in the genome chosen specified by the genomename argument. If FALSE, off-target calling will be skipped.
- *annotateoffs*: If TRUE, the function will provide annotations for the off-targets called using the genome annotation file specified by the gtfname argument. If FALSE, off-target annotation will be skipped.
- *getsgRNAdata*(x): This command is used to retrieve the data on the generated sgRNA sequences, where x is the raw data generated by *sgRNA_design*().
- *getofftargetdata*(x): This command is used to retrieve the additional off-target data, where x is the raw data generated by *sgRNA_design*().

crispRdesignR makes use of the R packages vtreat^9^, gbm^11^, Bsgenome^12^, BioStrings^14^, shiny^15^, and stringr^16^. Sequence homology features are calculated based on the gRNA interaction screen reported in Thyme *et. al*. ^17^.

## Results

The *crispRdesignR* tool is built entirely in the R programming language, utilizing various packages to assist with different aspects of the program (see Materials and Methods). The program can be run on the command line or through a graphical user interface (GUI). Guide RNAs are designed based on a 23 base pair sequence from a user-input DNA sequence or FASTA file that ends with the PAM. The only hard limitation on DNA regions that can be used as guide RNA is the presence of the PAM site, 5’NGG3’ in the case of spCas9, the most commonly used Cas9 enzyme. In order to effectively provide a score for the experimentally-supported scoring method used in *crispRdesignR*, flanking sequence is also collected; this flanking sequence includes the four base pairs before the 5’ end of the sgRNA and three base pairs after the 3’ end of the PAM sequence. In total, a region of 30 bases pairs is collected for each possible sgRNA. The R package searches for sgRNAs from the input and returns a table listing candidate sgRNAs and their sequence features, and optionally returns annotated off-target information in a user-chosen genome (Figure 1). The GC content of each target sequence is calculated excluding the PAM site, as the GC content of the PAM does not affect binding to the target region^3^. The self-complementarity score provided by *crispRdesignR* includes possible regions of self-complementarity within both the sgRNA target sequence and the region on the sgRNA backbone that is prone to forming hairpins. Homopolymers are detected by searching for strings of 4 or more consecutive base pairs.

**Figure 1.**
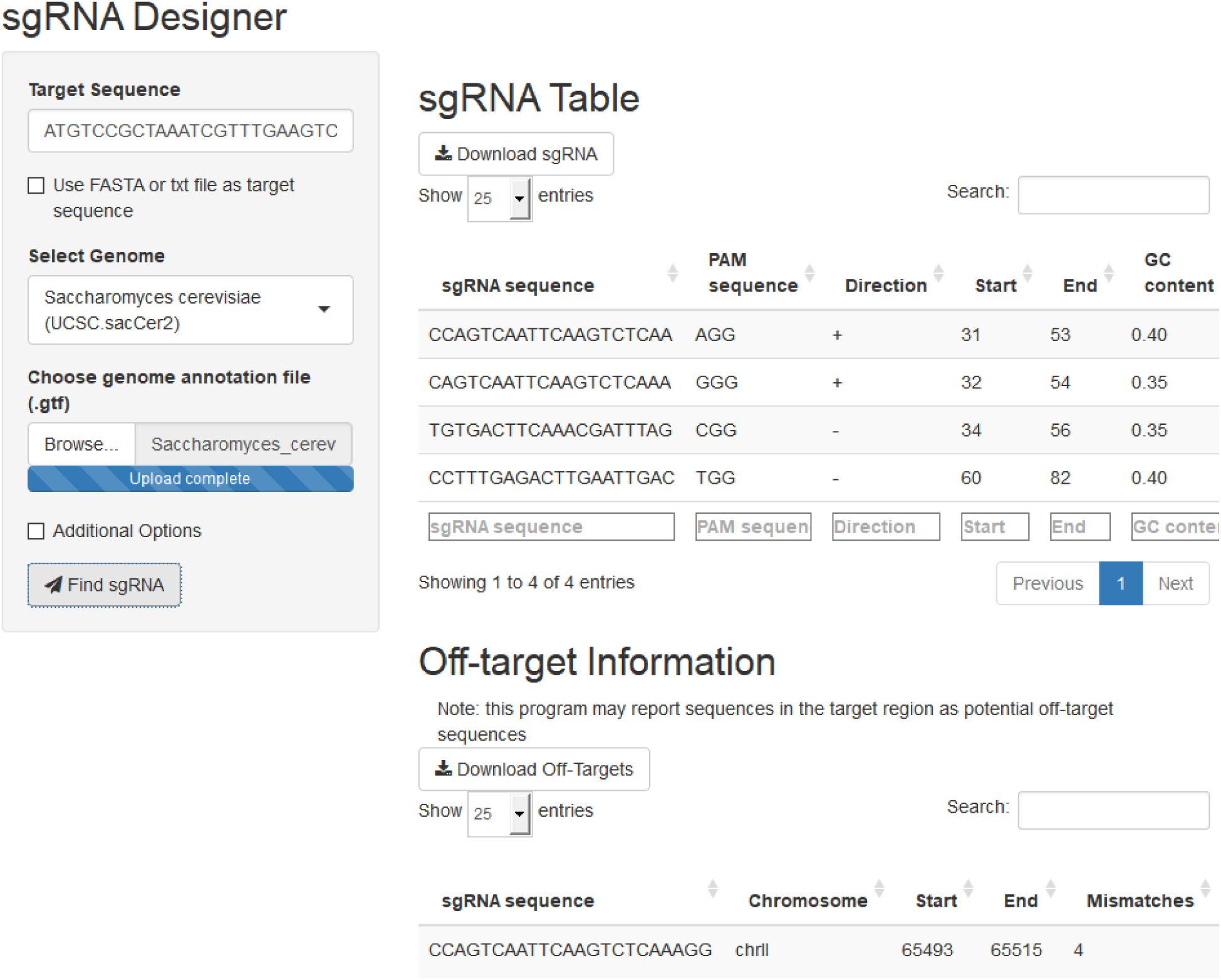
A screen capture from the *crispRdesignR* GUI demonstrating the target sequence, genome selection, and genome annotation file inputs. Partial sgRNA results and off-target annotations are also shown.

### Featurization

*crispRdesignR* has adopted the efficiency scoring method developed in Doench *et al*. (2016), employing a gradient boosted regression model trained on the FC and RES data set used in Doench Rule Set 2. In accordance with the Doench Rule Set 2, our model accounts for the presence of position-dependent single nucleotides, position-dependent dinucleotides, single nucleotide count, dinucleotide count, GC count, nucleotides that bookend the PAM sequence, and thermodynamic features of the target sequence plus context region (30-mer). The presence of specific nucleotides at certain positions in an sgRNA target site can influence the activity of that site. *crispRdesignR* considers the single and dinucleotides at each position and converts them into features that the machine learning model uses to predict activity.

To find off-target hits for the sgRNA, the genome from a user-selected species is loaded into the program through the Bsgenome^12^ package in R, and each guide RNA is then searched through the genome for up to 4 mismatches. Once a complete list of matching sequences with genomic locations has been collected, the program then cross-references the matching locations with gene information provided in a user-input gene annotation file (.gtf). If the sgRNA matches a position in a gene, *crispRdesignR* reports the gene name as well as whether the match lies in a coding region.

Running *crispRdesignR* will output two results tables (Figure 2). The first table contains the information on each individual sgRNA, including the sequence, PAM, location, direction relative to the target sequence, GC content, homopolymer presence, self-complementarity, off-target matches, and predicted efficiency score. The second table contains the information about each off-target match, including the original sgRNA, off-target sequence, chromosome, location, direction relative to the target sequence, number of mismatches, gene ID, gene name, type of DNA, and exon number. These tables can be sorted and searched through the GUI or downloaded as .csv files for further analysis. The location of the original sgRNA target sequence in the genome can be found in the off-target information section for identity verification. If no genome is provided or off-target searching is skipped, no data will be provided in the off-target matches column or the off-target information table.

**Figure 2.**
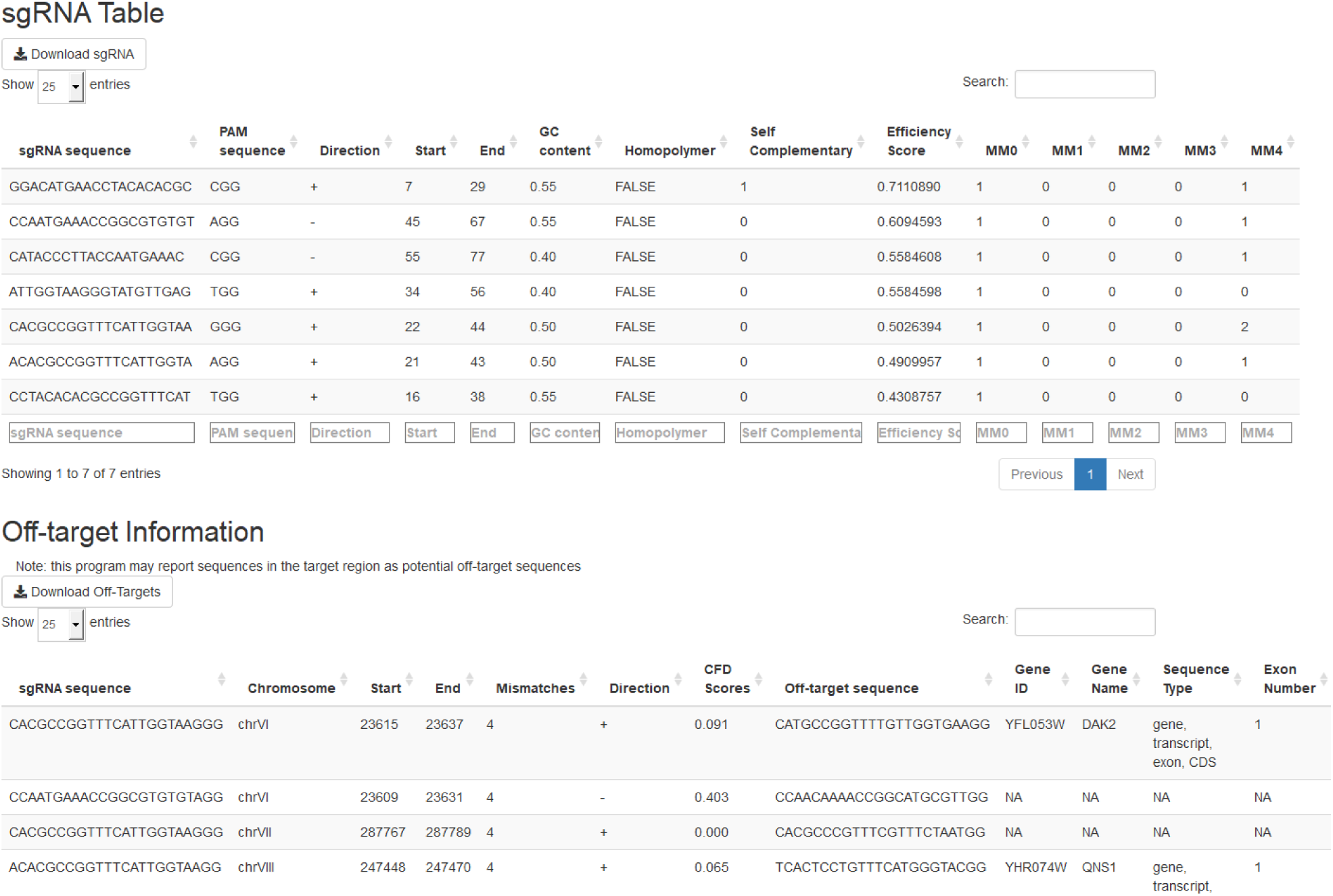
The output tables of *crispRdesignR* using a partial version of the DAK1 gene sequence, which is provided with the package download. Not all off-target matches are shown in the screenshot. Columns in the sgRNA table include sgRNA sequence, PAM, direction, start, end, GC content, presence of homopolymers, possible self-complementary sequences, efficiency score^3^, and number of matches in the user-provided genome with between 0 and 4 mismatches (MM). The Off-target information table includes the original sgRNA sequence, chromosome, start, end, number of mismatches, strand, CFD scores, matched sequence, gene ID, gene name, sequence type, and exon number.

### Benchmarking

Programs used to design sgRNA sequences often rely on predictive models but fail to report other sequence features that impact Cas9 enzymatic activity. In other cases, the information reported is calculated without excluding the PAM site, which is a recognition site for the protein and is not found in the sgRNA sequence. For example, CHOPCHOP v2^18,19^ is one of the few applications that will provide the GC content of each sgRNA sequence, but it provides the GC content of both the target sequence plus the PAM site, instead of the target site alone (however, this has been corrected in the newer version of CHOPCHOP (v3)^20^.

The *crispRdesignR* software excludes the PAM site from the sequence information reported and provides more sequence features to the user than other prominent free sgRNA design programs (Table 1). Its ability to search custom genomes and annotation files is essential when designing targets for non-model organisms and non-standard cell types. The ability to use customized PAMs in *crispRdesignR* permits the design of sgRNAs for uncommon Cas9 proteins. Another R-based program, CRISPRseek^21^, also allows users to design sgRNA in custom genomes with non-standard PAMs, but lacks a GUI and does not report several important sequence features such as hairpins, GC content, and homopolymers.

**Table 1.** Feature comparisons between several prominent free sgRNA design programs CHOPCHOP v2^18,19^, CRISPR Design^3^, CRISPRseek^21^, CRISPOR^22^, and GuideScan^23^. Features reported include whether all targets that match the PAM are output (All targets), the scoring method from Doench^3^, Moreno-Mateos^4^, or customizable), self-complementarity through hairpin detection, GC content, homopolymer filtering, the maximum number of mismatches permitted between the guide sequence and reference, the available PAM sequence, and whether off-target sequences are reported and annotated.

| Software name | CHOPCHOP v2<sup>18,19</sup> | CRISPR Design<sup>3</sup> | CRISPRseek<sup>21</sup> | CRISPOR<sup>22</sup> | GuideScan<sup>23</sup> | crispRdesignR |
| --- | --- | --- | --- | --- | --- | --- |
| Providing entity | Harvard | Broad Institute | UMASS Medical | Tefor | MSKCC | UML |
| All targets | Yes | No | Yes | Yes | Yes | Yes |
| Scoring method | Customizable | Doench | Doench | Doench & M.-Mateos | Doench | Doench |
| Hairpins | Yes | No | No | No | No | Yes |
| GC content | Yes | No | No | No | No | Yes |
| Homopolymers | No | No | No | No | No | Yes |
| Max no. of mismatches | 3 | 4 | 4 | 4 | 3 | 4 |
| PAM | Customizable | NGG, NNGRR | Customizable | Customizable | NGG, TTTN | Customizable |
| Off-target Annotation | No | Limited | Yes | Yes | No | Yes |

### Speed Comparisons

*crispRdesignR* has relatively fast runtimes to discover sgRNA sequences compared to other tools, although using custom genomes that are not pre-indexed leads to increased runtimes when choosing to call and annotate off-targets (Table 2). Most other web-based programs have pre-indexed genomes for fast off-target calling, but indexing can take several hours to perform and as such is not always ideal for users uploading custom genomes or for few queries. On a desktop with 3.4 GHz CPU and 8.00 GB RAM, the run time for a 128 bp sequence (“DAK1 short”, provided with the program) in *S. cerevisiae* averages out to 8 seconds in *crispRdesignR* when calling off-targets (3 seconds without off-target calling) compared to 7 seconds in *CRISPOR*^22^ and 5 seconds in *CHOPCHOP* v2^19^. GuideScan^23^ has some of the shortest runtimes when genomic coordinates are known beforehand and provided (2-3 seconds in *H. sapiens* and *S. cerevisiae*), but the web application can take over a minute if provided a FASTA file when searching the human genome. *crispRdesignR* and *CRISPRseek*^23^ are comparable in terms of speed, with *crispRdesignR* gaining a speed advantage when searching smaller genomes and *CRISPRseek* gaining an advantage in larger genomes. When performing off-target searches in the human genome, each additional sgRNA generated by *crispRdesignR* will add about 1 minute of run time. To reduce run-time when searching for off-targets, it is recommended that users keep DNA query sequences under 250 bases pairs when searching against a genome containing over a billion base pairs.

**Table 2.** Runtime comparisons for example sequences in each program analyzed. Run times (minutes:seconds) were averaged over three trials on a desktop PC with 3.4 GHz CPU and 8.00 GB RAM. Some programs offered a limited list of available genomes that prevented analysis (indicated by N/A). The DAK1 short example sequence can be found on the *crispRdesignR* github site; it is 128 bp long and generates 13 target sequences, with 35 off-targets. The DAK1 sequence contains 1780 bp and generates 170 target sequences, with 495 off-targets. The MYBPC3 deletion sequence contains 57 bp and generates 6 target sequences, with 2,219 off-targets. The Partial ADRB1 sequence contains 70 bp and generates 11 target sequences, with 9,200 off-targets.

| Test Sequence | Genome | CHOP-CHOP<sup>19</sup> | CRISPR Design<sup>3</sup> | CRISPRseek<sup>23</sup> | CRISPOR<sup>22</sup> | GuideScan<sup>23</sup> | crispRdesignR (no off-targets) | crispRdesignR (with off-target calling) |
| --- | --- | --- | --- | --- | --- | --- | --- | --- |
| DAK1 short | *S. cerevisiae* (yeast) | 0:05 | N/A | 2:10 | 0:07 | 0:02 | 0:03 | 0:08 |
| DAK1 | *S. cerevisiae* (yeast) | 0:18 | N/A | 4:24 | 0:19 | 0:02 | 0:14 | 1:47 |
| MYBPC3 deletion | *H. sapiens* (human) | 0:06 | 0:15 | 6:50 | 0:10 | 0:03 | 0:03 | 7:36 |
| Partial ADRB1 | *H. sapiens* (human) | 0:34 | 0:26 | 14:35 | 0:15 | 0:03 | 0:05 | 15:42 |

## Discussion

When utilizing other web-based sgRNA design programs, a user is often limited by a list of preinstalled genomes. *crispRdesignR* sets itself apart by allowing the user to import a custom genome and/or genome annotation file to search for sgRNAs and off-target effects. Allowing custom genomes and providing extensive target sequence information makes *crispRdesignR* particularly useful when working with non-model organisms, non-standard cell types and uncommon PAMs. The *crispRdesignR* software provides comprehensive sequence features to the user that are often omitted from other prominent free sgRNA design programs. The complete sequence feature information provided by *crispRdesignR* is very well-suited to applications where efficiency scores are of limited use. When using efficiency scoring methods with conditions that they have not been trained on (for example different organisms, cell types, and PAMs), the efficiency predictions will be less accurate. However, the predictive power of the model may not be completely lost if efficiency scoring methods are used in addition to known effects of various sequence features on activity to eliminate inactive sgRNA^3^.

The open source nature of *crispRdesignR* allows user to build on the features of the software for their specific uses. The gradient boosted regression model that *crispRdesignR* uses for efficiency scoring can be trained on other experimental data sets that contain the sgRNA sequence plus context (30-mer) and guide rankings assigned scores between 0 and 1. This allows for user-generated efficiency scoring models trained on data relevant to that user’s needs. However, for this to be a strongly predictive model, activity data must be available and normalized for thousands of sgRNA sequences in that relevant context^3^. The accessibility of the output tables as .csv files generated by *crispRdesignR* also allow a user to easily isolate the sgRNA sequences and run them through other scoring applications that are more appropriate for a specific application but that lack the sequence features, off-target annotation, or genome customization of *crispRdesignR*.

The flexibility and detail that is provided by the robust off-target annotation system used by *crispRdesignR* currently limits the speed of the program. While other programs may allow a user to index genomes for quicker searching, the process of indexing a custom genome can be hardware intensive and overall slower than a few searches on an unindexed genome for off-targets, particularly for design applications in a small target region. For applications that require sgRNA design in a large target region (over 1000 base pairs) within a large genome (over 1 billion base pairs), the user can turn off off-target calling in *crispRdesignR* to prevent long run times. Although web-based programs that access pre-indexed genomes offer superior speed, we show that they often report less sequence feature information, fewer off-targets, and they are limited to the genomes that can be searched to a pre-defined list.

Another R package, CRISPRseek^23^, uses similar methods of efficiency scoring and off-target calling, allowing for searching custom genomes and annotation files. However, it lacks the graphical user interface and several sequence features provided by *crispRdesignR*. The two programs both take longer to run than many of their web-based counterparts due to the ability to use non-indexed genomes, although *crispRdesignR* has a speed advantage when searching smaller genomes while *CRISPRseek* is faster when searching larger genomes. Although both programs use the same efficiency scoring method, *CRISPRseek* requires the user to add python packages in order to obtain the scores based on Doench Rule Set 2^3^. *crispRdesignR* is able to provide scores based on Rule Set 2 completely within R. Each program contains exclusive features that the other lacks that may be useful in different settings. For example, *CRISPRseek* has the ability to filter sgRNA based on restriction enzyme cutting sites, while *crispRdesignR* detects possible self-complementary sgRNA sequences.

The R package *crispRdesignR* sets itself apart by allowing the user to import a custom genome and/or genome annotation file to search for sgRNAs and off-target effects, while providing extensive target sequence information and the option of an accessible GUI. These unique features make *crispRdesignR* particularly useful for non-bioinformaticians working with non-model organisms, non-standard cell types, and uncommon PAMs. Accessible source code further adds to the versatility of *crispRdesignR* and lends itself to integration with different analysis pipelines and efficiency scoring methods as future technological improvements are made.

## Data Availability

The source code and example data for the *crispRdesignR* package is available at: https://github.com/dylanbeeber/crispRdesignR.

## Acknowledgements

The authors thank Evelyn Schwager and two anonymous reviewers for critical suggestions and feedback on the manuscript.

## Contributions

D.B. conceived the project. D.B. and F.C. designed and structured the R package. D.B. wrote the code for *crispRdesignR* and performed the analyses. D.B. and F.C. wrote the manuscript.

## References

1. Doudna JA, Charpentier E. The new frontier of genome engineering with CRISPR-Cas9. Science. 2014; 346.

2. Hsu P, Lander ES, Zhang F. Development and Applications of CRISPR-Cas9 for Genome Engineering. Cell. 2014; 157: 1262–1278.

3. Doench J, Fusi N, Sullender M, Hegde M, et al. Optimized sgRNA design to maximize activity and minimize off-target effects of CRISPR-Cas9. Nat. Biotechnol. 2016; 34: 185–195.

4. Moreno-Mateos M, Vejnar CE, Beaudoin JD, Fernandez JP, Mis EK, et al. CRISPRscan: designing highly efficient sgRNAs for CRISPR-Cas9 targeting in vivo. Nat. Methods. 2015; 12: 982–988.

5. Bolukbasi M, Gupta A, Wolfe S. Creating and evaluating accurate CRISPR-Cas9 scalpels for genomic surgery. Nat. Methods. 2016; 13: 41–50.

6. Haeussler M, Schoenig K, Eckert H, Eschstruth A, Mianne J, et al. Evaluation of off-target and on-target scoring algorithms and integration into the guide RNA selection tool CRISPOR. Genome Biol. 2016; 17(148).

7. Xie S, Shen B, Zhang C, Huang Z, Zhang Y. sgRNAcas9: A Software Package for Designing CRISPR sgRNA and Evaluating Potential Off-Target Cleavage Sites. PLoS One. 2014; 9(6).

8. Ma X. A robust CRISPR/Cas9 system for convenient high-efficiency multiplex genome editing in monocot and dicot plants. Mol. Plant. 2015; 8: 1274–1284.

9. Mount J, Zumel N. vtreat: A Statistically Sound ‘data.frame’ Processor/Conditioner. R package version 1.3.1; 2018.

10. Allawi H, SantaLucia, J. Thermodynamics and NMR of internal G.T mismatches in DNA. Biochemistry. 1997; 36(34).

11. Greenwell B, Boehmke B, Cunningham J. gbm: Generalized Boosted Regression Models. R package version 2.1.4; 2018.

12. Pagès H. BSgenome: Software infrastructure for efficient representation of full genomes and their SNPs. R package version 1.46.0; 2017.

13. R Core Team. R: A language and environment for statistical computing. R Foundation for Statistical Computing, Vienna, Austria; 2018.

14. Pagès H, Aboyoun P, Gentleman R, DebRoy S. Biostrings: Efficient manipulation of biological strings. R package version 2.46.0; 2017.

15. Chang W, Cheng J, Allaire JJ, Xie Yihui, McPherson J. Shiny: Web Application Framework for R. R package version 1.0.5; 2017.

16. Wickham H. stringr: Simple, Consistent Wrappers for Common String Operations. R package version 1.3.1; 2018.

17. Thyme S, Akhmetova, Montague TG, Valen E, Shier AF. Internal guide RNA interactions interfere with Cas9-mediated cleavage. Nat Commun. 2016; 7(11750).

18. Montague T, Cruz JM, Gagnon JA, Church GM, Valen E. CHOPCHOP: a CRISPR/Cas9 and TALEN web tool for genome editing. Nucleic Acids Res. 2014; 42: W401–W407.

19. Labun L, Montague TG, Gagnon JA, Thyme SB, Valen E. CHOPCHOP v2: a web tool for the next generation of CRISPR genome engineering. Nucleic Acids Res. 2016; 44: W272–W276.

20. Labun L, Montague TG, Krause M, Torres Cleuren YN, Tjeldnes H, Valen E. CHOPCHOP v3: expanding the CRISPR web toolbox beyond genome editing. Nucleic Acids Res. 2019; 47: W171–W174.

21. Zhu L, Holmes B, Aronin N, Brodsky M. CRISPRseek: a bioconductor package to identify target-specific guide RNAs for CRISPR-Cas9 genome-editing systems. PloS One. 2014; 9(9).

22. Concordet JP and Haeussler M. CRISPOR: intuitive guide selection for CRISPR/Cas9 genome editing experiments and screens. Nucleic Acids Res. 2018; 46: W242–W245.

23. Perez A, Pritykin Y, Vidigal J, Chhangawala S, Zamparo L, et al. GuideScan software for improved single and paired CRISPR guide RNA design. Nature Biotechnol. 2017; 35: 347–349.

